# Identification of a microRNA with a mutation in the loop structure in the silkworm Bombyx mori

**DOI:** 10.64898/2026.03.24.714027

**Authors:** Mayuko Harada, Midori Tabara, Kazunori Kuriyama, Katsuhiko Ito, Hidemasa Bono, Takuma Sakamoto, Miho Nakano, Toshiyuki Fukuhara, Atsushi Toyoda, Asao Fujiyama, Hiroko Tabunoki

## Abstract

MicroRNAs (miRNAs) play essential roles in the posttranscriptional regulation of gene expression in organisms. In the process of synthesizing mature miRNAs from miRNA precursors, the miRNA precursors are cleaved via Dicer at their loop structure, after which the miRNA precursors become mature and regulate transcription. However, the consequences of altering the loop sequence are not fully understood. The silkworm *Bombyx mori* is a lepidopteran insect with many genetic strains. We identified a mutant of the miRNA *miR-3260* whose the part of the loop structure was lacking in a silkworm strain with translucent larval skin. Here, we aimed to analyze the role of wild-type *miR-3260* and the influence of the mutation of the loop structure in *B. mori.* First, we identified the genomic region responsible for the translucent larval skin phenotype and determined that the mutated *miR-3260* nucleotide sequences. Then, we predicted the binding partners of wild-type *miR-3260* using the RNA hybrid tool and found two juvenile hormone (JH)-related genes as targets of wild-type *miR-3260*. Next, we assessed the relationships between *miR-3260* and JH and found that *miR-3260* was highly expressed in the Corpora allata and its expression responded to JH treatment. Meanwhile, *miR-3260* mimic and inhibitor did not induce the typical phenotypes associated with JH in *B. mori*. Then, we compared the dicing products from wild-type and mutant *miR-3260* precursors and observed that neither form underwent Dicer-mediated cleavage when the loop structure was altered. These results suggest that loop mutations in the *miR-3260* precursor may not influence dicing activity, consistent with the lack of observable phenotypic effects.

## Introduction

The silkworm *Bombyx mori* is a lepidopteran insect that has long been used to produce silk fiber. Advancements in genetic research have facilitated the breeding of silkworms that can produce high-quality silk, resist disease, and construct large cocoons, and silkworm races with favorable traits have been collected from around the world [1]. In this process, various phenotypes have been identified, and the silkworm races with these traits have been maintained as a genetic resource for use in the future [1]. The phenotypes of *B. mori* have been linked to causative genes via linkage maps. Currently, 561 races are maintained at the National Bioresource Project (NBRP) KAIKO [2].

*B. mori* has been used as a lepidopteran model insect in genetic, physiological, and pharmacological research [3]. The genome sequence of *B. mori* has been elucidated [4], and it has been reported that microRNAs (miRNAs) contribute to the regulation of gene expression in this organism [5,6].

miRNAs are short (18–25 bases), untranslated, single-stranded RNAs that play important roles in the posttranscriptional regulation of gene expression [7,8]. They bind to the 3’ untranslated region (3’UTR) of target gene mRNAs with incomplete complementarity and suppress the translation of the target gene [9]. miRNAs form a stem‒loop precursor when transcribed in the nucleus [10], are then cleaved by Dicer outside the nucleus [9,11], and undergo several modifications to become mature miRNAs [12]. The 2–8-base sequence at the 5’ end of mature miRNAs is called the seed sequence and binds to the target mRNA with perfect complementarity [13,14]. Each miRNA can target multiple different mRNAs, and the target mRNAs are degraded by specific miRNAs to control gene expression in the body [9].

Therefore, miRNAs posttranscriptionally regulate processes, such as metabolism, embryonic development, differentiation, disease response, and immunity, across organisms [12–16]. As of 2024, 38,589 hairpin precursors and 48,860 mature miRNAs from 271 organisms have been registered and published in the miRbase database [17].

Since the discovery of miRNAs, research into their biosynthetic pathways has been actively conducted. In particular, in the process of synthesizing mature miRNAs from miRNA precursors, the miRNA precursors are cleaved in a Dicer-dependent manner. It has been reported that Dicer recognizes the nucleotide (nt) sequence and secondary structure of the miRNA precursor at the cleavage site [18–22]. miRNA precursors have a loop structure [23,24], but this structure does not exist in mature miRNAs [25–27]. It has not been fully elucidated whether the absence of a correct nucleotide sequence in the loop structure is necessary for Dicer to recognize the loop structure [22,26,27].

We identified a mutant form of *miR-3260* lacking part of the loop structure in a silkworm mutant strain, o751 (*op* mutant), which displays spontaneous translucency during the larval stage, high pupal-stage mortality, and male infertility (Silkworm Genetic Resource Database: https://shigen.nig.ac.jp/silkwormbase/strain_detail.jsp). In this study, we aimed to analyse the role of wild-type *miR-3260* and the influence of the mutation of the loop structure in *B. mori*.

## Methods

### Silkworm strains

The wild-type (+*^op^*/+*^op^*) and the *op* mutant (*op*/*op*) o751 strains used were supplied by Kyushu University in the National Bio Resource project. The C108 and p50 strains were maintained by Tokyo University of Agriculture and Technology. For positional cloning, F_2_ progeny from the cross F_1_ (o75 × C108) female × F_1_ (o75 × C108) male were used. All silkworm larvae were reared on mulberry leaves at 25–28°C. The *B. mori* strain N4 used in this study was supplied by the University of Tokyo. Silkworm larvae were reared on the artificial diet silk mate 2S (Nosan, Tsukuba, Japan). Insects were maintained at 25°C with a 16-h light/8-h dark cycle.

### Cell culture

A silkworm cell line, BmN cells (RCB0457), was obtained from the Riken Cell Bank (Riken Tsukuba, Japan). BmN cells were maintained at 25°C in TC-100 medium (Thermo Fisher, Waltham, Massachusetts, USA) supplemented with 10% fetal bovine serum and an antibiotic-antimycotic mixture (Invitrogen, Carlsbad, CA).

### Positional cloning

For positional cloning, SNP and PCR markers were generated at various positions on chromosome 23, and the markers that were polymorphic between the parents were used for linkage analysis of 259 F_2_ larvae. The sequences of primers used in this study are shown in S2 Table. The candidate genes in the region narrowed by positional cloning were annotated using Silkbase [28].

### Preparation of genomic DNA and polymerase chain reaction (PCR) analysis

Genomic DNA was prepared using silk glands from fifth-instar larvae of the o-751 strain, employing DNAzol Reagent (Cosmo Bio) according to the manufacturer’s protocol. PCR was performed with Ex Taq DNA Polymerase (Takara Bio) using the primer sets listed in S3 Table, which were designed using the novel *Bombyx* draft genome sequence [29]. The PCR conditions were as follows: initial denaturation at 94°C for 2 min; 40 cycles of denaturation at 94°C for 15 s, annealing at 60°C for 15 s, and extension at 72°C for 1 min; and a final incubation step at 72°C for 4 min.

### Identification of the miRNA mutation

Genomic DNA was extracted from silk glands of the wild-type or *op* mutant from the o751 strain (n = 2 each) using a QIAprep Spin Miniprep Kit (QIAGEN, Venlo, Netherlands). The extracted genomic DNA was sheared to an average size of 400 bp using a Covaris Acoustic Solubilizer S220 (Covaris Inc., MA. USA) for paired-end library preparation. The sequencing libraries were constructed using the TruSeq DNA PCR-Free Sample Prep Kit (Illumina, CA, USA) according to the manufacturer’s instructions and sequenced on the HiSeq 2500 system (Illumina, CA, USA) with a read length of 2 × 150 bp. Based on the locus identified by positional cloning, we sought to differentiate the nucleotide sequences from those of the standard strain p50. We subsequently examined the target gene or miRNA, including the differentiation at the locus utilizing RNA sequencing (RNA-seq) data processed with HISAT2 (v2.0.1-beta) [30] and Stringtie (v1.2.2) [31]. *B. mori* miRNA data were obtained from miRbase [17]. By using the ultrafast sequence search tool GGGenome [32], those sequences were remapped to the newest genome assembly at that time. Only one miRNA (*bmo-mir-3260*) was mapped to the locus identified by positional cloning.

### Purification of miRNAs and total RNA from silkworm tissue

Whole bodies were collected and pooled for each developmental stage: eggs (n = 10), first-instar larvae (1L, n = 3), second-instar larvae (2L, n = 3), third-instar larvae (3L, n = 3), fourth-instar larvae (4L, n = 3), fifth-instar larvae (5L, n = 3), and pupae (n = 3). The corpora allata was dissected from day-4 fifth-instar larvae of the N4 *B. mori* strain (n = 10) and pooled into a single sample. Each sample was treated with QIAzol Lysis Reagent from the miRNeasy Mini Kit (Qiagen) or lysis buffer from the Pure Link RNA Mini Kit (Life Technologies) and then stored at −80°C until use. miRNA was purified via the miRNeasy Mini Kit (Qiagen) and RNeasy MinElute Cleanup Kit (QIAGEN) according to the supplier’s manual. Total RNA was purified via a Pure Link RNA Mini Kit (Life Technologies) according to the supplier’s manual.

### Synthesis of the cDNA library using purified miRNA and mRNA

A total of 200 μg of purified miRNA from each sample was used to synthesize the cDNA library via the TaqMan™ MicroRNA Reverse Transcription Kit (Thermo Fisher, Waltham, Massachusetts, USA). These cDNA libraries were stored at −30°C until use. One microgram of purified total RNA was treated with DNase I (Thermo Fisher), and cDNA was synthesized using the PrimeScript^TM^ 1st strand cDNA Synthesis Kit (Takara Bio Co. Ltd., Tokyo Japan) according to the supplier’s manual.

### Confirmation of the miRNA sequence

To confirm the mir-3260 sequence, DNA purified from the silk glands of 4-day-old fifth-instar larvae of *B. mori* o751 and the p50 strain (n = 2 each) was used as a template to amplify the miRNA sequence using KOD Plus Neo (TOYOBO, Tokyo, Japan), and the primer pair 5’-ATGCAAGATAGAAGAAGTAG-3’ and 5’-TCAACTCACGAAAGTAATC-3’ was designed on the basis of the genome sequence. The amplified PCR products were electrophoresed on a 1% agarose gel, and bands of the objective nucleotide length were excised and extracted using the QIAquick Gel Extraction Kit (50) (QIAGEN, Venlo, Netherlands). 3’A was added to both ends of the extracted PCR products using 10×A-attachement Mix (Toyobo Co., Ltd., Tokyo, Japan) and then ligated into the pCR 2.1-TOPO vector (Life Technologies, Waltham, Massachusetts, USA). The vector with the miRNA sequence ligated was transformed into ECOS^TM^ Competent *E. coli* XL1-Blue (NIPPON GENE, Tokyo, Japan). The transformed *E. coli* cells were then inoculated onto LB agar medium supplemented with 100 μg/mL ampicillin and incubated at 37°C for 16 h. The emerged colonies were examined by direct colony PCR with M13 primers (S3 Table) to determine whether the target *bmo-mir-3260* sequence had been introduced. The clone of interest was cultured with shaking at 37°C for 16 h in liquid LB medium supplemented with 100 μg/mL ampicillin. The culture was centrifuged at 3,500 rpm for 20 min to collect the bacterial cells, and the plasmid DNA was purified using a QIAprep Spin Miniprep Kit (QIAGEN, Venlo, Netherlands). The nucleotide sequence of the resulting plasmid DNA was confirmed by the Sanger method using a 3730xl DNA analyzer (Life Technologies, Waltham, Massachusetts, USA).

### Reverse transcription quantitative PCR for miRNA

For the template, 2 μL of cDNA was used. TaqMan™ Fast Advanced Master Mix (Thermo Fisher, Waltham, Massachusetts, USA) and the *miR-3260* primers (Thermo Fisher, TaqMan miRNA Assay, Assay ID: 241990_mat) were mixed with each miRNA cDNA library, and quantitative RT‒PCR was performed using the Step One Plus Real‒Time PCR system (Applied Biosystems) according to the manufacturer’s instructions. The relative expression levels of the miRNAs were calculated using the delta‒delta Ct method from the Ct values obtained by quantitative RT‒PCR. U6 (Thermo Fisher, TaqMan miRNA Assay, Assay ID: 001973) was used as an endogenous control. miRNA expression levels in whole bodies are presented as RQ values. Whole-body miRNA expression was normalized to first-instar larvae (RQ = 1), whereas corpora allata expression was normalized to fifth-instar, whole-body larval samples (RQ = 1). Error bars indicate RQ minimum and maximum values based on standard deviation. All assays were performed in technical triplicate.

### Reverse transcription quantitative PCR for total RNA

For the template, 2 μL of cDNA was employed. RT-qPCR was also performed on the cDNA libraries synthesized from total RNA using primers for each gene and the KAPA SYBR Fast qPCR Kit (NIPPON Genetics) according to the manufacturer’s instructions. The relative mRNA expression level for each gene was calculated using the delta‒delta Ct method from the Ct values obtained by quantitative RT‒PCR. Ribosomal protein 49 (Rp49) was used as an endogenous control (S3 Table). mRNA expression levels in each sample are presented as RQ values, normalized to control samples (RQ = 1). Error bars reflect RQ minimum and maximum levels based on standard deviation. All assays were performed in triplicate.

### Prediction of miRNA secondary structure and target mRNA binding

The secondary structures based on wild-type or mutant *miR-3260* nucleotide sequences were predicted by RNAfold WebServer [33]. We searched for mRNA sequences that bind to miRNAs using transcriptomic data from the standard silkworm strain p50T and the mutant strain o-751 (PRJDB19213). The method for RNA-seq analysis is in the Supporting Information. First, we identified homologous miRNA sequences from miRbase, confirmed their mature sequences, and then obtained seed sequences. *miR-3260* is located on chromosome 23, adjacent to cluster of JH-related genes (Chr23; 9916751-10377814), according to the silkworm gene database. Coordinated expression of neighboring genes has been documented in various species [30]. Therefore, we speculated that *miR-3260* expression may be involved in regulating nearby JH-related genes. To investigate this, we retrieved data on the relevant gene clusters from KAIKObase [31]. Next, we extracted the 3’ untranslated region (3’UTR) sequence, which is the sequence following the stop codon that is the target of the miRNA, from the transcriptome data of the o-751 strain larvae on the 4th day of the 5th instar. Using these data and the JH-related gene cluster sequence, we used RNAhybrid [32] to search for mRNAs with sequences complementary to the seed sequence (Figs. 1–3). As conditions for binding to the target sequence, the mfe, which indicates the ease of binding, was set to −20 or less, and the seed sequence was allowed to have a G:U wobble.

**Fig. 1.**
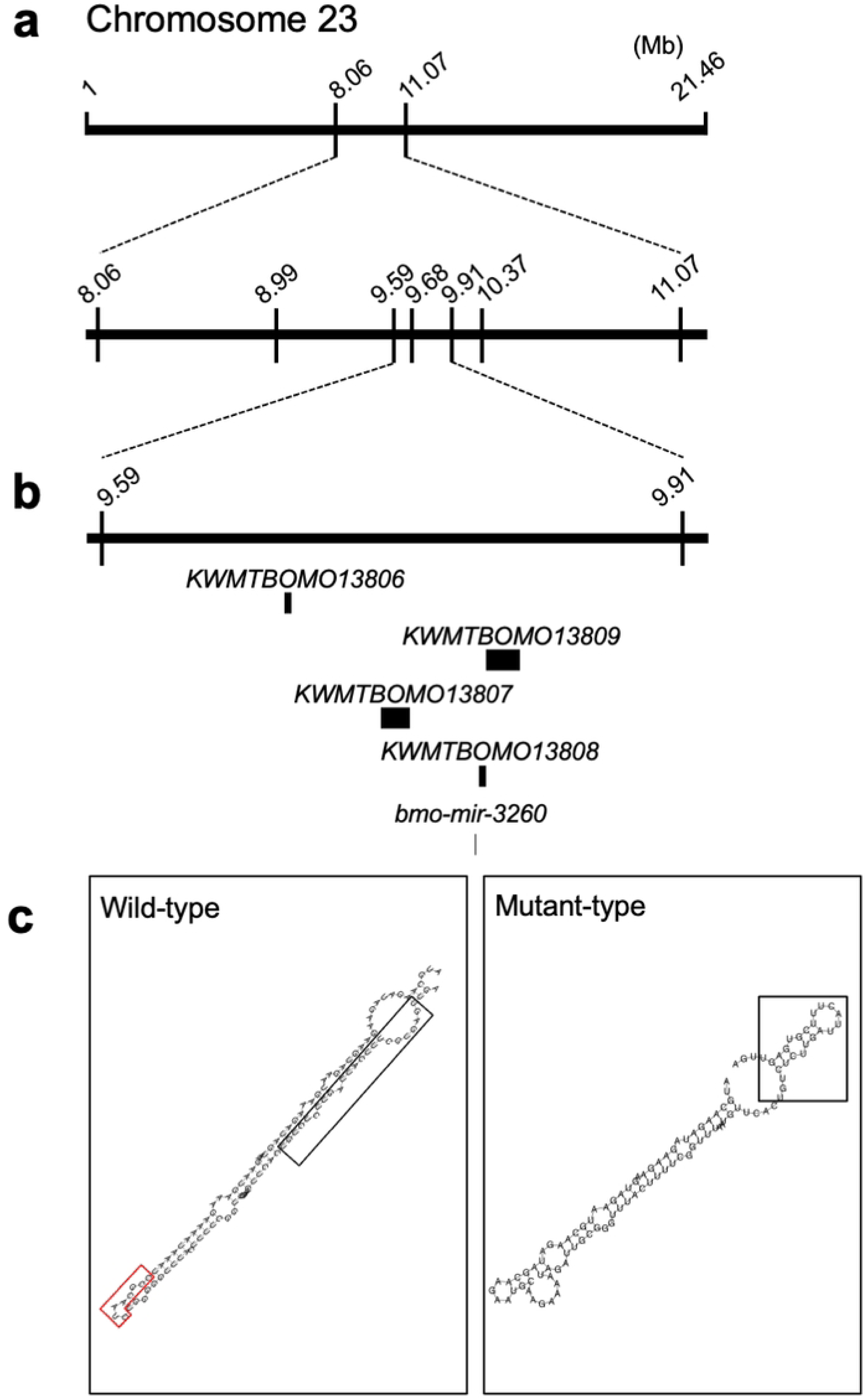
Positional cloning of the *op* gene on chromosome 23. (**a**) Positional cloning of chromosome 23. Each number indicates the chromosome position of the marker listed in S2 Table. The *op* locus was located within a region approximately 0.32 Mb in length between 9,598,832 and 9,916,751 bp on chromosome 23. (**b**) Predicted genes within the narrowed region. This region contains four predicted genes, namely, *KWMTBOMO13806*, *KWMTBOMO13807*, *KWMTBOMO13808*, and *KWMTBOMO13809*, and one miRNA, namely, *bmo-mir-3260*. (**c**) Prediction of the miRNA secondary structure. The secondary structure was predicted on the basis of the miR-3260 nucleotide sequence via RNAfold WebServer. Wild-type indicates wild-type *mir-3260*, and mutant indicates m*mir-3260*. The black box indicates the mature sequence region, and the red box indicates missing nucleotide sequences in the *miR-3260* mutant.

**Fig. 2.**
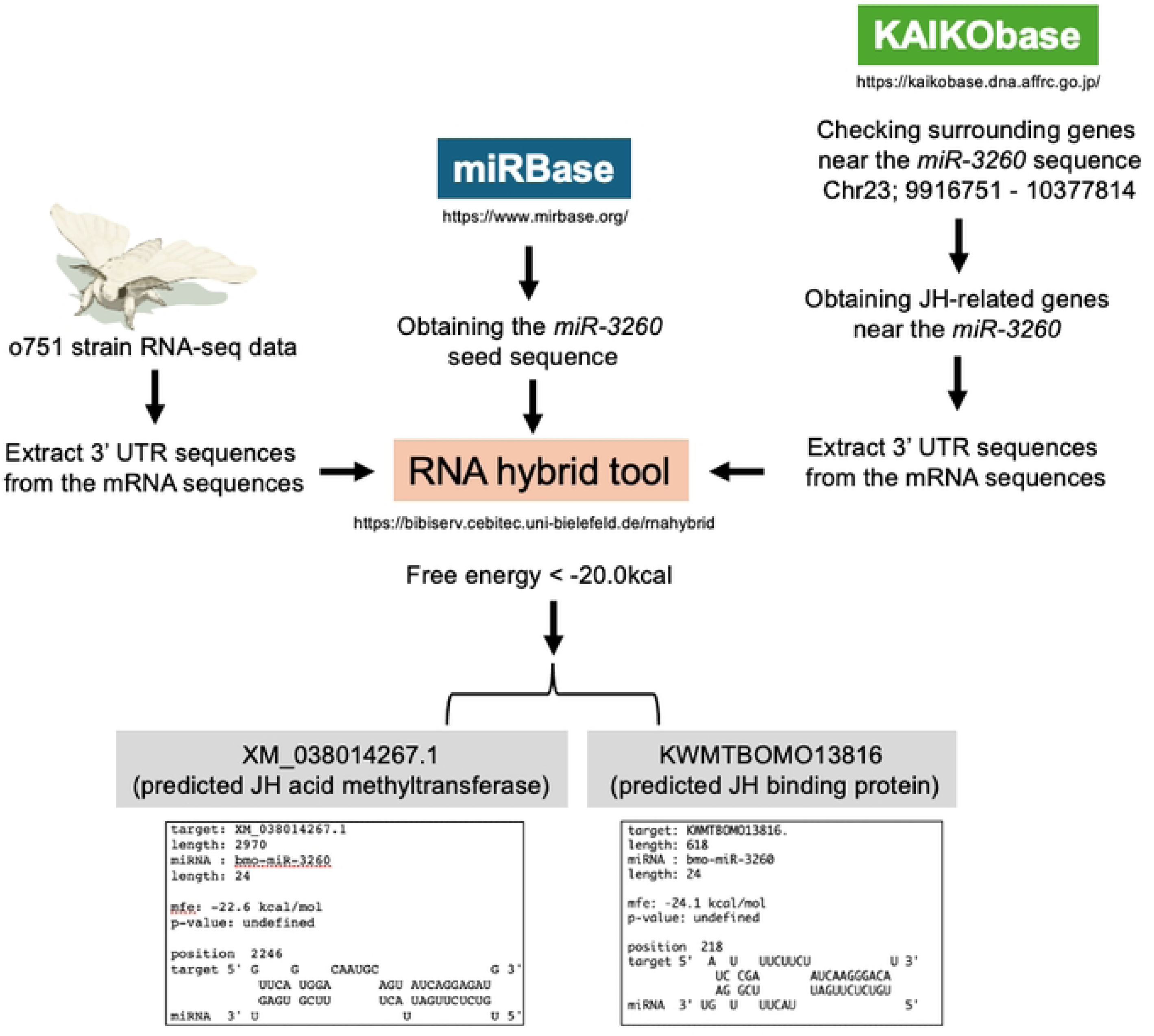
The process for the Prediction of target mRNA binding. mRNA sequences that bind to *miR-3260* were predicted using RNAhybrid. The minimum free energy value (mfe), which indicates the ease of binding, was set to −20 or less, and the seed sequence was allowed to have a G:U wobble.

**Fig. 3.**
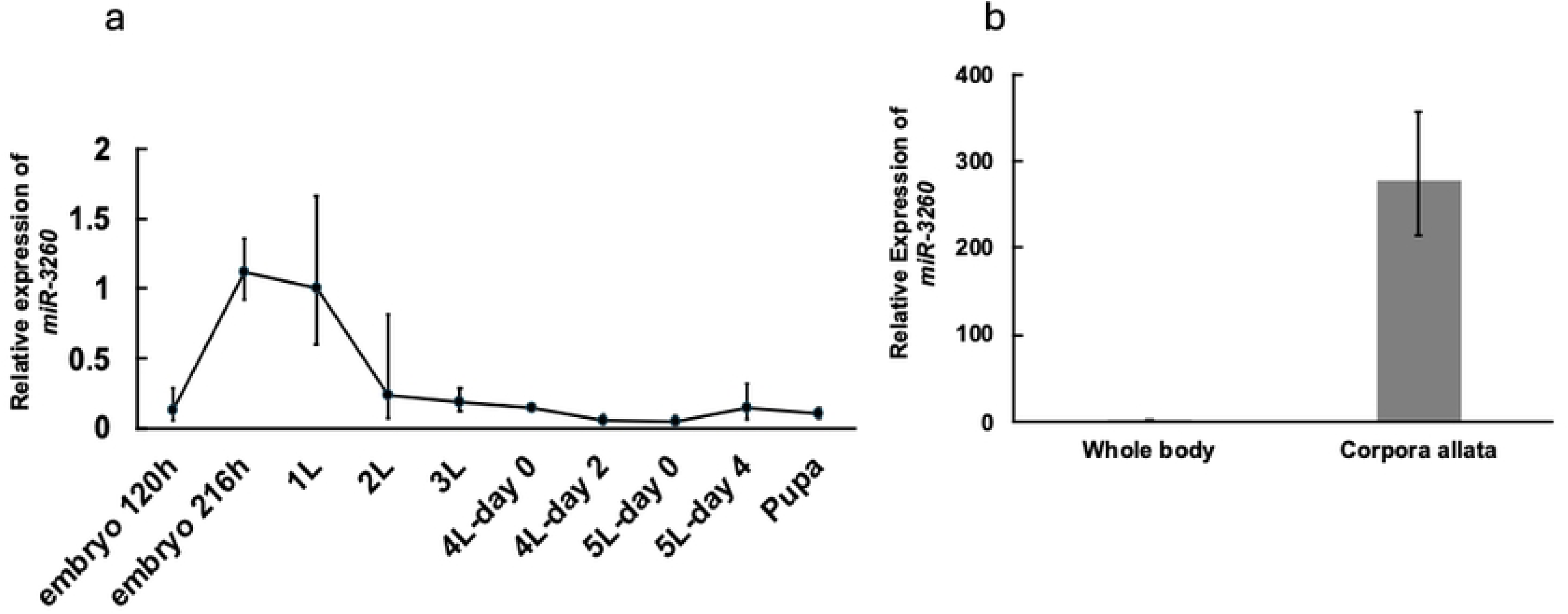
*miR-3260* expression in the developmental stage and corpora allata. (**a**) *miR-3260* expression during development. Embryo 120 h and 216 h indicate 120 and 216 h after oviposition, respectively; 1L, 2L, and 3L indicate first-, second-, and third-instar larvae, respectively; 4L-day 0 and 4L-day 2 indicate days 0 day and 2 of the fourth-instar larvae, respectively; 5L-day 0 and 5L-day 2 indicate days 0 day and 2 of the fifth-instar larvae, respectively. Whole-body miRNA expression levels are presented as RQ values, representing relative expression levels normalized against first-instar larvae samples (RQ = 1). Error bars represent RQ minimum and maximum levels based on standard deviation. U6 was used as an endogenous control. All sample sets were assayed in triplicate as technical replicates. (**b**) *miR-3260* expression in the corpora allata. The miRNA expression levels in the corpora allata are presented as RQ values, which represent the relative expression levels calculated by assuming the value for whole bodies of 4-day fifth-instar larval samples as 1. Error bars indicate RQ minimum and maximum levels based on standard deviation. U6 was used as an endogenous control. All sample sets were assayed in triplicate as technical replicates.

### Examination of miR-3260 expression by JH treatment

BmN cells (2 × 10^6^) were grown on 6-well Falcon plates (BD Biosciences, Franklin Lakes, NJ). JH III was purchased from Cayman Chemical (Michigan, USA) and dissolved in 10% isopropanol to prepare a 10-mg/ml stock solution. The stock solution was diluted with 1x insect PBS to 100 μg/mL. In total, 50 μL of the diluted JH solution was added to each well with BmN cells (n = 3), yielding a final JH concentration of 9.4 μM per well. For controls, a 10% isopropanol solution was diluted 1:100 with 1x insect PBS, and 50 μL of this solution was added to each well with BmN cells (n = 3). JH-treated BmN cells were cultured at 25°C for 24 and 48 h, harvested, pooled by treatment group, and subjected to miRNA purification and RT-qPCR.

### Injection of miR-3260 mimic or inhibitor into B. mori eggs

Female and male adults of the N4 strain of *B. mori* were mated, and their eggs were used for injection within 4 hours after laying. Using an electric manipulator and a microinjector, holes were made in the eggs with a tungsten needle, and the egg liquid was attached to the surface of the eggshell. Then, 5 nL of the 100mM of *miR-3260* mimic (Thermo Fisher, product ID: MC18121, S4 Table) or inhibitor (Thermo Fisher, product ID: MH18121, S4 Table) was injected into the egg liquid with a glass capillary, and it was confirmed that the egg liquid had returned to the inside of the egg. The hole was sealed with instant adhesive. Water was used as a control. Eggs were incubated at 25°C until hatching (n = 48) or until reaching the second-instar larval stage (n = 5). Whole-body samples of second-instar larvae were used for *Jhamt* and *KB13816* mRNA expression analysis.

### Dicing assay

Dicing (Dicer) assays were performed according to the methods of Tabara et al. [34]. BmN cells were cultured in T25 flasks, and dissected fat bodies were sonicated on ice in extraction buffer [20 mM Tris-HCl (pH 7.5), containing 4 mM MgCl_2_, 5 mM DTT, 1 mM PMSF, 1 μg/mL leupeptin, and 1 μg/mL pepstatin A]. BmN cell and fat body lysates were centrifuged twice at 13,000 × *g* and 4°C for 15 min, and supernatants were collected with protein concentrations adjusted to 0.1 mg/mL. Subsequently, 1 μL of ^32^P end-labeled dsRNA or miRNA precursor single-stranded RNA (ssRNA) (*pre-bmo-let7, pre-bmo-*m*let7, pre-miR-3260*, and *pre-*m*miR-3260*; Agilent Technologies, Palo Alto, CA, USA; S5 Table) was incubated with 15 μL of BmN or fat body lysate containing BmDicer-1 and BmDicer-2 proteins and 4 μL of 5× dicing buffer [18 mM HEPES-KOH (pH 7.4), with 7.5 mM DTT, 3.3 mM magnesium acetate, 100 mM potassium acetate, 0.25% glycerol, and 1 mM ATP] at 25°C for 30 min. Postincubation, fractions were separated via 15% denaturing PAGE and analyzed using autoradiography. Cleaved RNAs were visualized as band intensities using a Typhoon FLA 7000 image analyzer (GE Healthcare Japan) after 24 h, and original autoradiography images were processed in high-contrast mode using ImageQuant TL (GE Healthcare; S2 Fig.).

### Statistical analysis

Statistical significance was assessed in R (ver4.3.1) using Fisher’s exact test with multiple comparisons. Holm’s method is used to correct the familywise error rate. *P* < 0.05 was considered statistically significant.

## Results

### Positional cloning of the op locus and searching for the op phenotype

To identify the genomic region responsible for the *op* phenotype, we performed positional cloning using the *Bombyx* draft genome sequence [28,29]. Through genotyping using 259 F_2_ individuals, we delimited the locus to an approximately 320-kb-long region on chromosome 23 (Fig. 1a and S1 Table). A search against silkBase [28] revealed that this region contained four predicted genes, namely, *KWMTBOMO13806*, *KWMTBOMO13807*, *KWMTBOMO13808*, and *KWMTBOMO13809*, and one miRNA, namely, *bmo-mir-3260* (Fig. 1b). We subsequently compared the sequences of the region containing these candidate genes with the genome sequence of the standard strain p50. We identified a *bmo-mir-3260* mutant lacking an 8-nucleotide sequence in the o-751 strain silkworm, and the deletion was located in the loop structure in *bmo-mir-3260* (Fig. 1c). We refer to the wild-type *bmo-mir-3260* as *miR-3260* and the *bmo-mir-3260* lacking the loop structure as m*miR-3260*.

### Determination of the mir-3260 sequence

To determine the nucleotide sequence of *miR-3260*, we amplified the target miRNA via polymerase chain reaction (PCR) using DNA extracted from the silk glands of p50 or o-751 strains (+/+ or *op*/*op*, respectively) as templates. The 110-base sequences were obtained from the o-751 strain. We then confirmed whether there was an m*miR-3260* sequence in the *op*/*op*, +/*op*, and +/+ o-751 strains and found that m*miR-3260* was conserved in the o-751 strain but not in the p50 strain. Sequence analysis revealed that *mmiR-3260* is eight nucleotides shorter than wild-type *miR-3260* and differs at five nucleotide positions (S1 Fig.). The m*miR-3260* nucleotide sequence was submitted to the International Nucleotide Sequence Database (DNA Data Bank of Japan (DDBJ)) under accession number LC858578.

### Prediction of miR-3260 target genes

By using the *B. mori* transcriptome data, we searched for target mRNAs that bind to *miR-3260* (Fig. 2). Genes located close together on the same chromosome often exhibit coordinated expression, as demonstrated in humans, mice, rats, yeast, fruit flies, and worms [35]. The region near *mir-3260* (Chr23; 9916751-10377814) contained a juvenile hormone (JH)-related gene cluster. We speculated that *miR-3260* expression may be linked to the expression of these JH-related genes. Thus, we focused on the JH-related gene cluster close to *miR-3260* on Chr23 (9916751-10377814).

We obtained 3’UTR sequences of the JH-related genes near *miR-3260* from KAIKObase [36] and 3’UTR sequences from our RNA sequencing (RNA-seq) data and then analyzed their binding ability via the RNAhybrid tool [37]. On the basis of the results, *B. mori* juvenile hormone acid methyltransferase (Jhamt, XM_038014267.1) from our RNA-seq data, and *B. mori* juvenile hormone binding protein (JHBP) ce-0330 (*KWMTBOMO13816*, AB196703) from KAIKObase data were predicted as candidate genes that may bind to *miR-3260* (Fig. 2). Therefore, two JH-related genes were predicted to be binding partners of *miR-3260*.

### Examination of miR-3260 expression during the developmental stage and in the corpora allata

We analyzed *miR-3260* expression from the embryonic to pupal stages (Fig. 3a). Compared with the first-instar larval stage, *miR-3260* expression levels in embryos at 216 h after oviposition and in first-instar larvae were higher. Expression of *miR-3260* gradually decreased after the early larval developmental stages, such as in first- and second-instar larvae (Fig. 3a). We also examined *miR-3260* expression in the corpora allata, which synthesizes and secretes JH. The expression of *miR-3260* in the corpora allata of the 5th-instar larvae on the 4th day was also examined and was found to be significantly greater than that in the whole-body (Fig. 3b).

### Examination of miR-3260 expression in response to JH treatment in BmN cells

We further explored the relationship between *miR-3260* and JH by treating BmN cells with JH and measuring *miR-3260* expression levels at 24 and 48 h. We found that *miR-3260* expression increased approximately 2.5-fold compared with that in the control group at 24 h (Fig. 4a). The expression level of *miR-3260* also increased approximately 6-fold compared with that in the control group at 48 h (Fig. 4b). Thus, *miR-3260* responded to JH treatment in BmN cells.

**Fig. 4.**
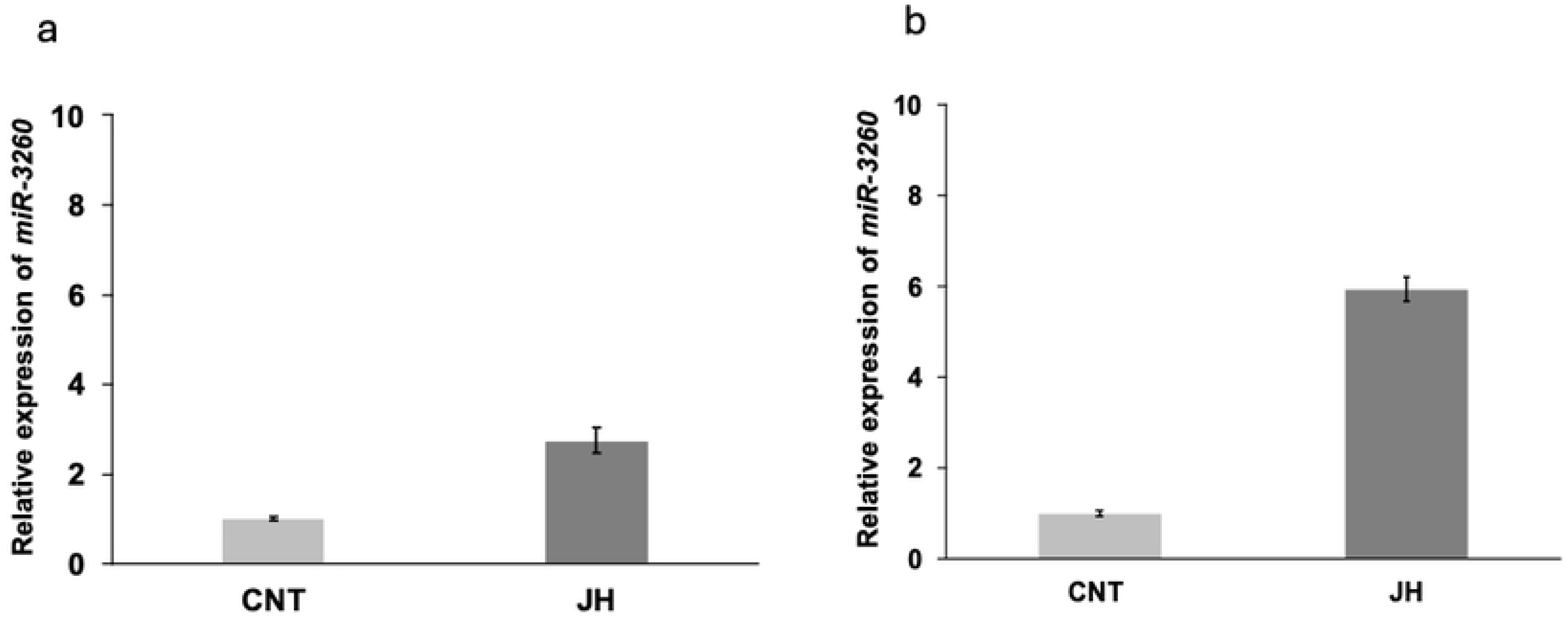
*miR-3260* expression in BmN cells treated with juvenile hormone for 24 h (a) or 48 h (b). The miRNA expression levels in the BmN cells are presented as RQ values, which represent the relative expression levels calculated assuming the value for the control samples as 1. Error bars represent RQ minimum and maximum levels based on standard deviation. U6 was used as an endogenous control. All sample sets were assayed in triplicate as technical replicates.

### Examination of JH-related gene expression during the developmental stage

We investigated the expression of *Jhamt* and *KWMTBOMO13816* mRNAs from the embryonic to the pupal developmental stages. *Jhamt* mRNA expression was highest in the embryo 120 h after oviposition (Fig. 5a), whereas *KWMTBOMO13816* mRNA expression was highest at 216 h after oviposition, followed by a sharp decline in the early larval stages (Fig. 5b).

**Fig. 5.**
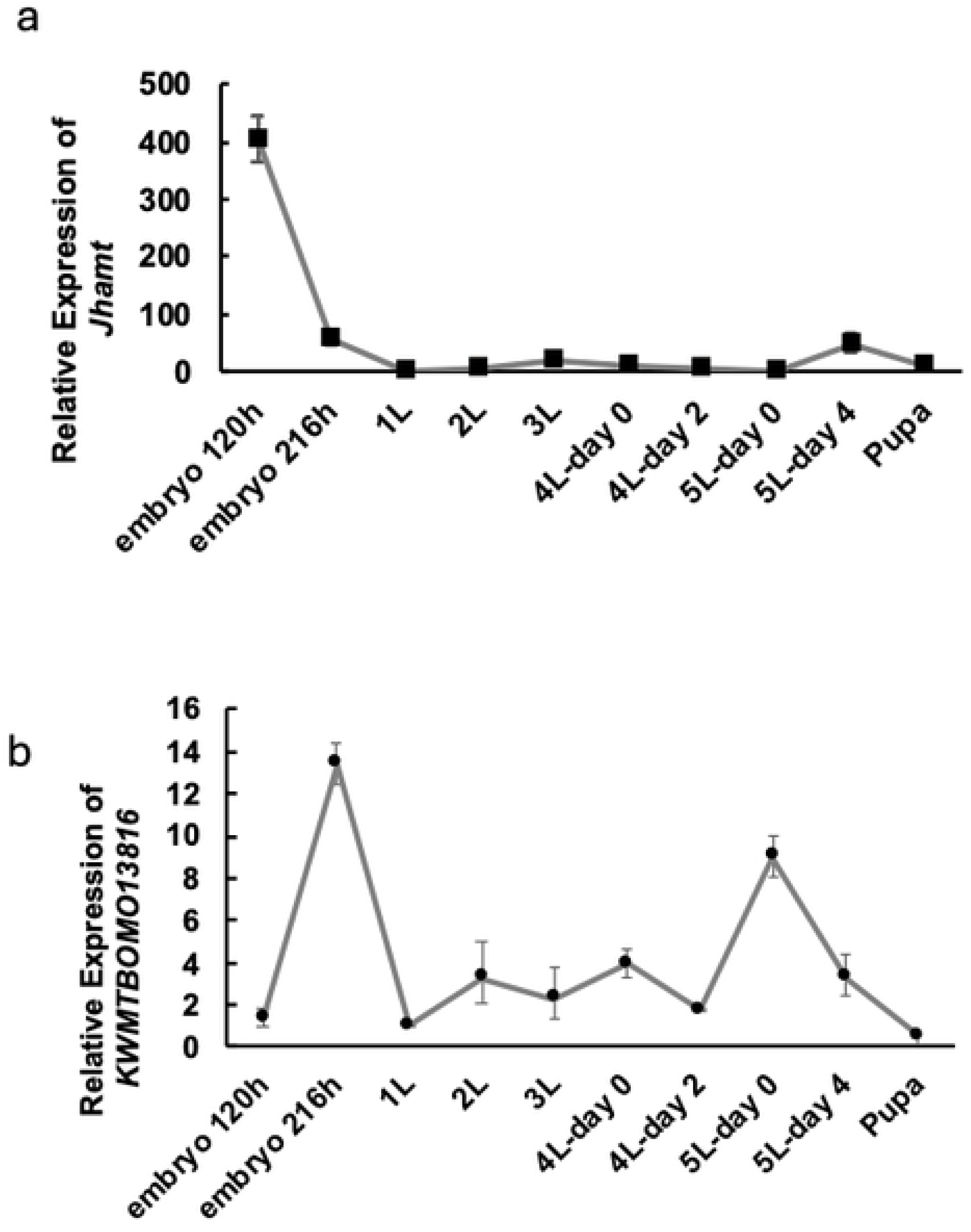
*Jhamt* and *KWMTBOMO13816* mRNA expression at the developmental stage. (**a**) *Jhamt* mRNA expression, (**b**) *KWMTBOMO13816* mRNA expression. Embryo 120 h indicates 120 hours after oviposition; Embryo 216 h indicates 216 hours after oviposition; 1L indicates first-instar larva; 2L indicates second-instar larva; 3L indicates third-instar larva; 4L-day 0 indicates day 0 of the fourth-instar larva; 4L-day 2 indicates day 2 of the fourth-instar larva; 5L-day 0 indicates day 0 of the fifth-instar larva. 5L-day 2 indicates day 2 of the fifth-instar larva. mRNA expression levels in the whole bodies are presented as RQ values, which represent the relative expression levels calculated by assuming the value for the first-instar larval samples as 1. Error bars indicate RQ minimum and maximum levels based on standard deviation. *Rp49* was used as an endogenous control.

### Examination of the phenotype by injecting a miR-3260 mimic or inhibitor

We assessed the phenotype and relationship between *miR-3260* and the expression of *Jhamt* and *KWMTBOMO13816* during embryogenesis by injecting a *miR-3260* mimic or inhibitor. The expression patterns of *miR-3260*, *Jhamt*, and *KWMTBOMO13816* suggested a possible relationship during embryonic and early larval development. However, injecting the *miR-3260* mimic (*P* = 0.3221) or inhibitor (*P* = 0.1071) at larval hatching did not affect hatching rate or larval appearance compared with the control (Fig. 6a). Subsequently, *miR-3260* mimic was injected into eggs, and *Jamt* and *KWMTBOMO13816* mRNA expression levels were measured following development to the second-instar larval stage (Fig. 6b and c). *Jhamt* mRNA was markedly upregulated following injection of either *miR-3260* mimic or inhibitor into eggs (Fig. 6b). In contrast, *KWMTBOMO13816* mRNA expression slightly decreased with *miR-3260* mimic injection but remained unchanged with inhibitor injection (Fig. 6c).

**Fig. 6.**
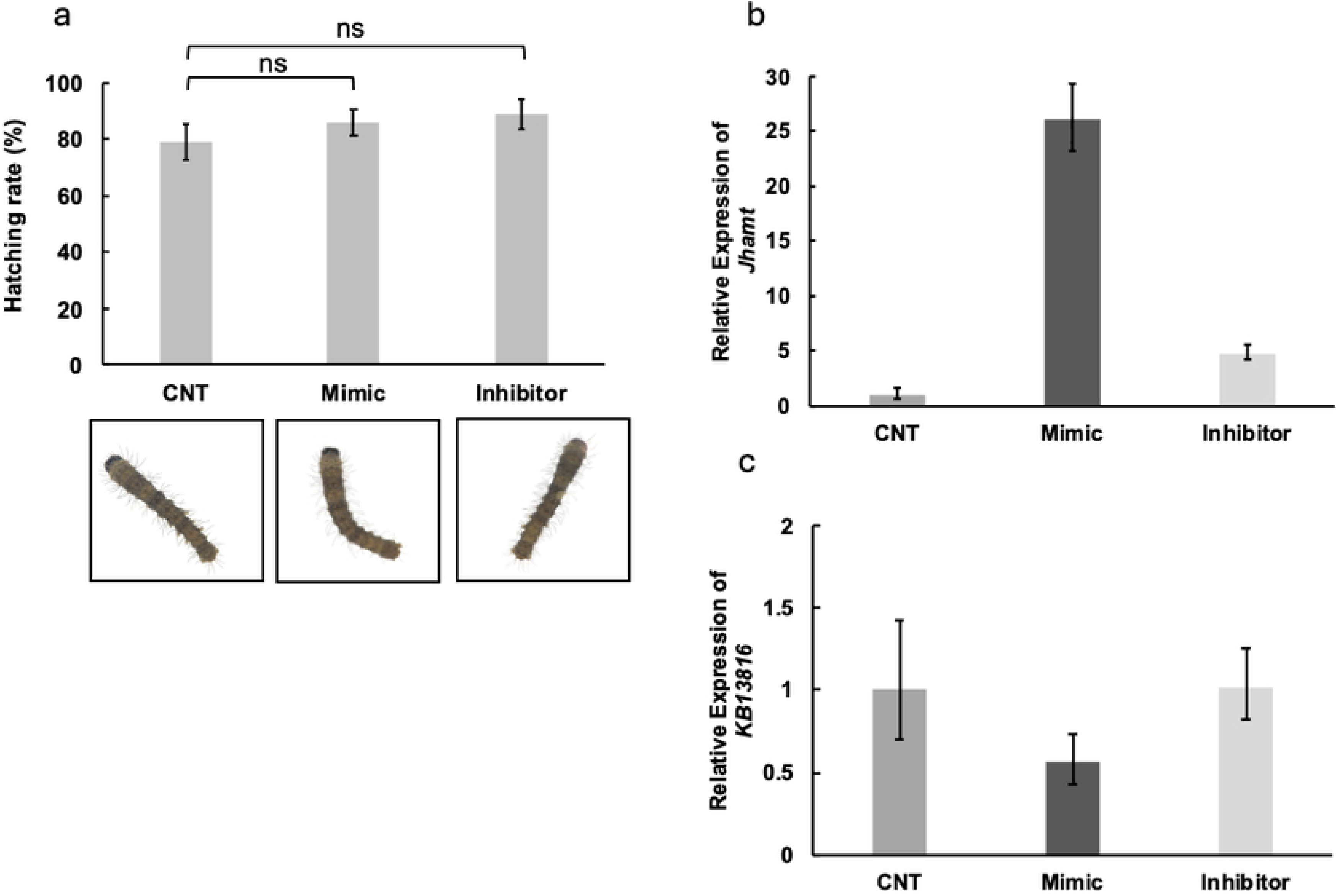
Examination of the effects of injecting the *miR-3260* mimic or inhibitor. (**a**) The upper graph shows the hatching rate; the lower panel shows the phenotype of the first-instar larva. ns indicates non significance. (**b**) *Jhamt* mRNA expression after injection of the *miR-3260* mimic or inhibitor. (**c**) *KWMTBOMO13816* mRNA expression after injection of the *miR-3260* mimic or inhibitor; mRNA expression levels in each sample are presented as RQ values, which represent the relative expression levels calculated assuming the value for the control samples as 1. Error bars represent RQ minimum and maximum levels based on standard deviation. Rp49 was used as an endogenous control.

### Examination of dicing activity toward pre-miR-3260 and pre-mmiR-3260

The stem‒loop structure in the miRNA precursor [9] is processed by Dicer [10,11]. This process plays a crucial role in the biosynthesis of mature miRNAs [12]. To examine how alterations in the stem‒loop structure affect the production of mature miRNA, we performed a dicing assay using cell-free crude extracts prepared from BmN cells and fat body. When we used a 50-nt perfectly matched double-stranded RNA (dsRNA) with blunt ends as a substrate, a 20-nt RNA cleaved by BmDicer-2 was detected (Fig. 7a and d, red asterisks), as reported previously [34]. The wild-type *bmo-let7* precursor (*pre-bmo-let7*) was cleaved into a 22 nt fragment (miRbase, MIMAT0015221) by putative BmDicer-1 activity (Fig. 7b and d, gray asterisks), but the mutant *bmo-let7* precursor *(pre-bmo-*m*let7*) was slightly cleaved (Fig. 7d, red asterisks). Mature *miR-3260* is 24 nt in length, according to the information in miRbase (MIMAT0015444). Although ^32^P-labeled *pre-miR-3260* appeared more degraded than *pre-mmiR3260*, neither precursor exhibited cleavage at the expected ∼24-nucleotide positions (Fig. 7c and d).

**Fig. 7.**
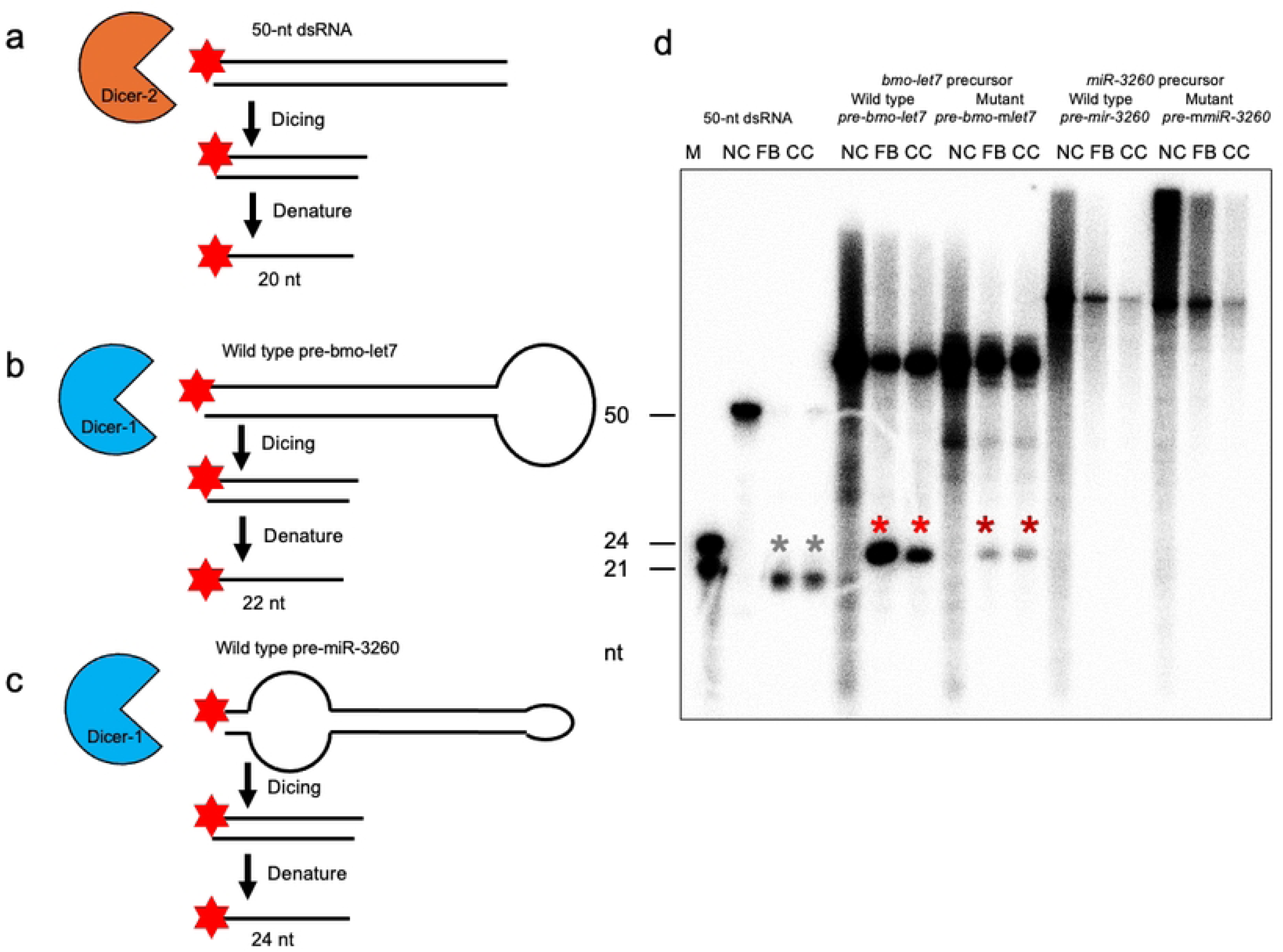
Comparison of dicing activity between *pre-miR-3260* and *pre-*m*miR-3260*. A dicing assay was performed using crude extracts prepared from BmN cells as enzyme fractions and ^32^P-end-labeled ssRNA or dsRNA as substrates. The cleaved RNAs were detected as band intensities with a Typhoon FLA 7000 image analyzer. Schematic diagram of the dicing assay used in this study: (**a**) 50-nt dsRNA, (**b**) wild-type bmo-let7 precursor, and (**c**) wild-type miR-3260 precursor. The red star indicates ^32^P. (**d**) The 50-nt dsRNA was used as a dicing assay experimental control. The bmo-let7 precursor (*pre-bmo-let7*) was used as a dicing assay positive control. The mutant *bmo-let7* precursor (*pre-bmo-*m*let7*) was used as a dicing assay negative control. Wild-type and mutant of the miR-3260 precursor. Lane M, RNA size markers of 50, 24, and 21 nt; NC, negative control (no-enzyme fraction); FB, fat body; CC, cultured BmN cells. Red or gray asterisks indicate diced products by BmDicer-1 and BmDicer-2, respectively.

## Discussion

In this study, we found that m*miR-3260* lacked some of the nucleotide sequences in the loop structure. First, we looked for *miR-3260* binding partners and identified *Jhamt* and *KWMTBOMO13816* as candidates. When a miRNA binds to a target mRNA, the binding affinity is indicated by the free energy value, and it has been reported that binding may occur when the minimum free energy (mfe) is −20.0 kcal/mol or less; the lower the free energy value is, the stronger the binding [37,38–40]. The mfe for *Jhamt* and *miR-3260* was −22.6, and that for *KWMTBOMO13816* and *miR-3260* was −24.1. Bioinformatics analysis suggested that *miR-3260* may bind to these target mRNAs.

*miR-3260* expression was markedly upregulated from 216 h after oviposition to the first-instar larval stage, coinciding with the period of new JH synthesis [41]. Additionally, *miR-3260* expression was significantly higher in the corpora allata of fifth-instar larvae on day 4. Furthermore, treatment of BmN cells with JH also upregulated *miR-3260* expression, and previous studies have reported sustained mRNA expression of *Jhamt* and *KWMTBOMO13816* expression after dorsal closure during embryogenesis [41]. These findings suggest that *Jhamt and KWMTBOMO13816* mRNA expression levels are correlated with *miR-3260* expression. *Jhamt* is the enzyme that methylates farnesoic acid, producing the final product JH [42]. *KWMTBOMO13816* is annotated as a JHBP, ce-0330, in the *B. mori* compound eye expressed sequence tag database [43]. Many JH-binding proteins are secreted into the haemolymph, bind with JH, and transport JH to the cells [43]. However, the *KWMTBOMO13816* amino acid sequence has been shown to lack secretion signal sequences, and *KWMTBOMO13816* is predicted to play a role in the cytosol [43]. We found that *miR-3260* showed peak expression from 216 h after oviposition to the first-instar larval stage, whereas *KWMTBOMO13816* expression declined significantly in first-instar larvae. The JH titer during embryonic developmental is estimated to increase from 96 h and remain elevated after hatching [41]. We speculated that *miR-3260* may regulate *KWMTBOMO13816* expression in the embryonic developmental stage.

Next, we assessed the phenotypes and relationships among *miR-3260*, *Jhamt*, and *KWMTBOMO13816* during embryogenesis. We used a *miR-3260* mimic and inhibitor.

miRNAs bind to transcripts with a complementary sequence in the 3’ region and control protein expression [7,8]. Thus, the *mir-3260* mimic or inhibitor was injected into *B. mori* eggs, and the phenotypes and mRNA expression of *Jhamt* and *KWMTBOMO13816* were examined. The hatching rates did not differ among the groups, and the *miR-3260* mimic or inhibitor did not affect the appearance of the first-instar larvae. Next, by injecting a *miR-3260* mimic into eggs, we assessed whether *miR-3260* inhibits the expression of *Jhamt* or *KWMTBOMO13816* at second-instar larvae. Injection of the *miR-3260* mimic significantly upregulated *Jhamt* mRNA expression while only slightly downregulating *KWMTBOMO13816*. Similarly, the *miR-3260* inhibitor upregulated *Jhamt* mRNA expression but did not affect *KWMTBOMO13816*. Therefore, the outcomes did not fully align with the predicted regulatory functions of the *miR-3260* mimic or inhibitor. Individuals lacking the ability to synthesize or receive JH have been confirmed to develop normally inside the eggshell but do not hatch in *B. mori* [41]. It has been reported that JH synthesis or knockout of the JH receptor is not affected during embryonic development, as no phenotype is observed [41]. Despite slightly reducing *KWMTBOMO13816* expression, *miR-3260* mimics or inhibitors did not produce clear phenotypic changes, indicating that a regulatory effect remains possible but may be limited.

As mentioned above, the stem‒loop structure in the miRNA precursor plays a crucial role in processing mature miRNAs. m*miR-3260* lacked an 8-nucleotide sequence in its loop structure. Dicer recognizes the 3’ overhanging end structure of the substrate and measures the distance from that structure to the single-stranded RNA region, and the binding ability decreases when the stem structure is long or when the loop structure is small [44]. Dicer cleaves pre-miRNAs into duplexes comprising a guide strand and a star strand, with the guide strand incorporated into Argonaute (AGO) and the star strand subsequently degraded [45]. In our dicing assay, Dicer recognition relied on the 5`′` phosphate site. Although wild-type *miR-3260* possesses a mature sequence in the 3′ terminal site, the assay system lacks AGO activity; thus, cleavage fragments may persist as miRNA duplexes. Despite ^32^P-labeled pre-miR-3260 showing some degradation, neither wild-type nor mutant miR-3260 generated fragments corresponding to the 24-nt sequence size. Even if minor effects occurred on target genes, such as *Jhamt* and *KB13816*, embryonic development in *B. mori* was unaffected by mimics or inhibitors, suggesting that *miR-3260* does not behave like a typical miRNA, and loop alterations alone are unlikely to change the phenotype.

We remain uncertain why wild-type *mir-3260* responds to JH and is highly expressed in the corpus allatum. However, qRT-PCR confirmed changes in its expression, implying a functional role, although the mechanisms underlying these expression patterns remain unresolved. Current experimental approaches are limited in their ability to clarify the function of miRNAs with unique loop structures. Future technological advances may provide further insights. Based on our results the stem‒loop structure of *miR-3260* may be resistant to canonical Dicer cleavage, suggesting that both wild-type and mutant *miR-3260* may not act as conventional miRNA.

## Conclusion

We analyzed wild-type *miR-3260* function and performed a dicing assay to impact the loop mutations. Neither form exhibited the canonical dicing pattern, predicted in miRbase, reinforcing the notion that *miR-3260* may function differently from typical miRNAs. Remaining questions include why *miR-3260* is highly expressed during embryogenesis and in the corpus allatum, and why it responds to JH. Future studies will investigate how loop mutations influence *miR-3260* function and explore the relationship between JH signaling and *miR-3260* during *B. mori* embryogenesis.

## Supporting information

**S1 Fig.** Alignment of wild type and mutant type of *miR-3260*. WT indicates wild type, and MT indicates mutant type. The multiple sequence alignment was performed using CLUSTAL O (1.2.4). The hyphens indicate a lack of nucleotide sequence. The black asterisks indicate the same nucleotide sequences between wild-type and mutant types of *miR-3260*. The gray asterisks indicate the different nucleotide sequences between wild-type and mutant types of *miR-3260*. The red asterisks indicate mature sequences.

**S2 Fig.** Original autoradiography image for Fig. 7 d. The original autoradiography image was processed using ImageQuant TL with High contrast mode.

**S1 Table.** Positional cloning of the *op* candidate gene.

**S2 Table.** Primers used for positional cloning.

**S3 Table.** Primers used for miRNA cloning and qRT‒PCR.

**S4 Table.** Nucleotide sequences used for the mimic and inhibitor of *B.mori miR-3260*.

**S5 Table.** Nucleotide sequences used for the chemical synthesis of pre-miRNAs and dsRNA.Fig.

## Acknowledgments

We thank Dr. Kikuo Iwabuchi, Tokyo University of Agriculture and Technology, for helpful discussion and Ms. Saori Matsumoto, Tokyo University of Agriculture and Technology, for technical assistance.

## Funding

This work was supported by JSPS KAKENHI grants 18H02212, 20K20571, and 23K17418 to HT and 22K14899 to TS. This work was also supported by the Center of Innovation for Bio-Digital Transformation (BioDX), an open innovation platform for industry–academia cocreation (COI-NEXT), Japan Science and Technology Agency (JST, JPMJPF2010), provided to H.B. and H.T. The ROIS-DS-JOINT grants 004RP2017 and 016RP2018 were provided to H.B. and H.T.

## Author contributions

Conceptualization: M.H., M.T., K.I., H.B., A.F., T.F., and H.T.

Methodology: M.H., M.T., K.K., K.I., H.B., T.S., T.F., A.T., A.F., and H.T.

Investigation: M.H., M.T., K.K., K.I., T.S., M.N., A.T., A.F., T.F., and H.T.

Visualization: M.H., M.T., K.K., K.I., H.B., T.S., T.F., and H.T.

Supervision: H.T.

Writing—original draft: M.H., K.I., T.S., T.F., and H.T.

Writing—review & editing: M.T., K.K., K.I., H.B., T.S., M.N., T.F., A.T., A.F.

## Data availability statement

The raw sequence data were deposited in DDBJ under the umbrella BioProjects PRJDB19213 (RNA-Seq reads) and PRJDB19532 (Illumina short reads of the genomes of the wild-type and *op* mutant).

The datasets used and/or analyzed during the current study available from the corresponding author on reasonable request.

## Additional information

Supporting information is available online.

## Competing interests

The authors declare no competing interests.

## Notes

### Competing Interest Statement

The authors have declared no competing interest.

